## Supplementary material for "Pan-cancer characterization of ncRNA synergistic competition uncovers potential carcinogenic biomarkers": S1 File

### Supplementary Methods

#### Predicting ceRNA networks in Pan-cancer

Three criteria (significant sharing of miRNAs, significantly positive correlation, and significantly sensitive correlation conditioning on shared miRNAs) are used to predict ceRNA network.

The first criterion uses a hypergeometric test to evaluate the significance of the sharing of miRNAs between ncRNA *a* and mRNA *b*. The significance *p*-value of the test is calculated as:

$$p_{ab} = 1 - \sum_{x_{ab}=0}^{L_{ab}-1} \frac{\binom{M_{ab}}{x_{ab}}\binom{N_{ab}-M_{ab}}{K_{ab}-x_{ab}}}{\binom{N_{ab}}{K_{ab}}} \qquad (1)$$

where $N_{ab}$ is the number of all miRNAs in Pan-cancer dataset, $M_{ab}$ and $K_{ab}$ represent the total numbers of miRNAs interacting with ncRNA *a* and mRNA *b* respectively, and $L_{ab}$ is the number of miRNAs shared by ncRNA *a* and mRNA *b*.

The second criterion requires that ncRNA *a* and mRNA *b* is positively correlated. The Pearson [1] method is utilized to assess whether the expression level of ncRNA *a* and the expression level of mRNA *b* are strongly and positively correlated at a significant level (e.g. 0.05). The correlation $cor_{ab}$ between ncRNA *a* and mRNA *b* is computed as:

$$cor_{ab} = cor(x, y) = \frac{\sum_{i=1}^{s}(x_i - \bar{x})(y_i - \bar{y})}{\sqrt{\sum_{i=1}^{s}(x_i - \bar{x})^2}\sqrt{\sum_{i=1}^{s}(y_i - \bar{y})^2}} \qquad (2)$$

where $x = (x_1, x_2, ..., x_s)$ and $y = (y_1, y_2, ..., y_s)$ denote the expression level of ncRNA *a* and mRNA *b* respectively, $\bar{x}$ and $\bar{y}$ represent the average expression level of ncRNA *a* and mRNA *b* respectively, and *s* is the number of samples for each tumor type. Correspondingly, the significance *p*-value $pc_{ab}$ is calculated as:

$$pc_{ab} = 2pt\left( cor_{ab}\sqrt{\frac{s-2}{1-(cor_{ab})^2}} \right) \tag{3}$$

where the *pt*() R function is used to calculate the probability of $cor_{ab}\sqrt{\frac{s-2}{1-(cor_{ab})^2}}$.

The third criterion is used to evaluate the influence of the shared miRNAs on ncRNA *a* and mRNA *b*. The sensitive correlation [2] is employed to calculate the influence of the miRNAs shared by ncRNA *a* and mRNA *b*, and the null model [3] is applied to evaluate the significance of the influence. The sensitive correlation $sc_{ab}$ is calculated as:

$$sc_{ab}=cor_{ab} - pcor_{ab} \tag{4}$$

where $pcor_{ab}$ is the partial correlation between ncRNA *a* and mRNA *b*, i.e. the correlation conditioning on their shared miRNAs, or the correlation between ncRNA *a* and mRNA *b* when the influence of the shared miRNAs is omitted. $pcor_{ab}$ is computed as:

$$\begin{aligned} pcor_{ab} &= cor(x,y\,|\,Z) = cor(x,y\,|\,(Z_1,Z_2,...,Z_m)) \\ &= \frac{cor(x,y\,|\,(Z_1,Z_2,...,Z_{m\text{-}1})) - cor(x,Z_m\,|\,(Z_1,Z_2,...,Z_{m-1}))cor(y,Z_m\,|\,(Z_1,Z_2,...,Z_{m-1}))}{\sqrt{1-cor(x,Z_m\,|\,(Z_1,Z_2,...,Z_{m-1}))^2}\sqrt{1-cor(y,Z_m\,|\,(Z_1,Z_2,...,Z_{m-1}))^2}} \end{aligned} \tag{5}$$

where $x$, $y$ and $Z$ denote the expression level of ncRNA *a*, mRNA *b*, and the shared *m* miRNAs respectively, $cor(x,y\,|\,(Z_1,Z_2,...,Z_m))$ denotes the partial correlation between *x* and *y* conditioning on $(Z_1,Z_2,...,Z_m)$, $cor(x,y\,|\,(Z_1,Z_2,...,Z_{m-1}))$ represents the partial correlation between *x* and *y* conditioning on $(Z_1,Z_2,...,Z_{m-1})$, $cor(x,Z_m\,|\,(Z_1,Z_2,...,Z_{m-1}))$ is the partial correlation between *x* and $Z_m$ conditioning on $(Z_1,Z_2,...,Z_{m-1})$, $cor(y,Z_m\,|\,(Z_1,Z_2,...,Z_{m-1}))$ is the partial correlation between *y* and $Z_m$ conditioning on $(Z_1,Z_2,...,Z_{m-1})$。

To evaluate the significance of $sc_{ab}$, the null model assumes that the shared miRNAs do not affect the correlation between ncRNA *a* and mRNA *b*, i.e. the sensitive correlation (the difference between $cor_{ab}$ and $pcor_{ab}$) between ncRNA *a* and mRNA *b* is 0. The number of datasets sampled is set to 1E+03 for the null model, and the pre-computed covariance matrices in SPONGE R package [3] are used to build the null model. Based on the constructed null model, we can infer the significance *p*-value of $sc_{ab}$.

In this work, a candidate ncRNA-mRNA pair with *p*-value < 0.05 for significant sharing of miRNAs (criterion 1), significantly positive correlation (criterion 2) and significantly sensitive correlation conditioning on shared miRNAs (criterion 3) is regarded as a ceRNA interaction. After integrating all of ceRNA interactions in each tumor type, we can predict ceRNA networks across 31 tumor types in Pan-cancer.

#### Multi-class classification metrics

In the R package utiml [4], we use 22 metrics (accuracy, average-precision, clp, coverage, F1, hamming-loss, macro-AUC, macro-F1, macro-precision, macro-recall, margin-loss, micro-AUC, micro-F1, micro-precision, micro-recall, mlp, one-error, precision, ranking-loss, recall, subset-accuracy, wlp) to evaluate the performance of multi-class classification. The metric accuracy is the percentage of correct predictions (both true positives and true negatives) among the retrieved samples. The metric average-precision is the average of the precisions of all possible recalls. The metric constant label problem (clp) is the inability of a multilabel classification (MLC) strategy to correctly predict the same class for all samples. The metric coverage computes the average number of classes that have to be included in the final prediction in order to cover all the true classes. The metric F1 is the harmonic mean of the precision and recall. The metric hamming-loss is the percenatge of the wrong classes to the retrieved classes. The metrics macro-AUC, macro-F1, macro-precision, macro-recall are the average of AUC, F1, precision and recall over all classes. The metric margin-loss is the loss that uses a margin to compare samples representations distances. The metrics micro-AUC, micro-F1, micro-precision, micro-recall are calculated by considering adding all the true positives, true negatives, false positives and false negatives for each class, and add them up to compute them. The metric missing label problem (mlp) is the inability of an MLC strategy to correctly predict a specific class. The metric one-error measures how many times the top ranked predicted class is not in the set of true classes of the sample. The metric precision (also called positive predictive value) is the percentage of relevant samples among the retrieved samples. The metric ranking-loss is an error measure that computes the averaged rate of class pairs that are inversely sorted. The metric recall (also known as sensitivity) is the percentage of relevant samples that were retrieved. The metric subset-accuracy is the percentage of samples that have all their classes classified correctly. The metric wrong label problem (wlp) is the inability of an MLC strategy to correctly predict one of the existing classes at least for one sample in the whole dataset.

#### Construction of the web-based SCOMdb resource

The web-based source was organized by MySQL (version 5.7.17-log) and queried using JavaServer Pages (JSP). The web interface was developed using HTML5 with JavaScript. All resources in SCOMdb were stored and managed using MySQL (version 5.7.17-log). The web interface was built in JSP. The data processing programs were written in Java (version 1.8.0_31), and the web services were built using Apache Tomcat. The SCOMdb source is freely available at www.comblab.cn/SCOMdb/.

#### Validation of synergistic competition ncRNAs as carcinogenic biomarkers

The manual curation of experimentally validated carcinogenic biomarkers is obtained from LncACTdb 3.0 [5]. We are only interested in the carcinogenic biomarkers associated with our 31 malignant tumors. We have found that none of carcinogenic biomarkers in Pheochromocytoma and Paraganglioma (PCPG) and Mesothelioma (MESO) are experimentally validated by the literature.

### Supplementary Figures

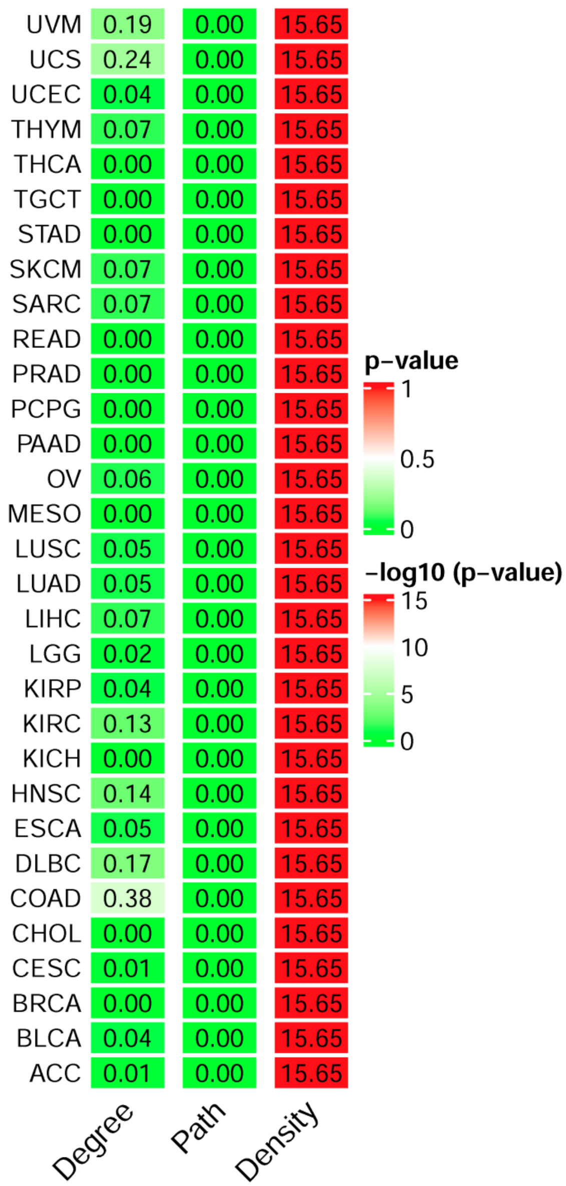


**Fig S1. The topological characteristics of ceRNA networks across the 31 malignant tumors.** The Degree column is the *p*-value of fitting power-law distribution. Small *p*-values (less than 0.05) indicate that the degree of ceRNA networks do not obey the power-law distribution. The Path and Density columns represent the –log10(*p*-value) of Student's *t*-test. Larger values in the Path column denote that the characteristic path length of ceRNA networks is significantly shorter than that of random networks, and larger values in the Density column indicate that the density of ceRNA networks is significantly higher than that of random networks.

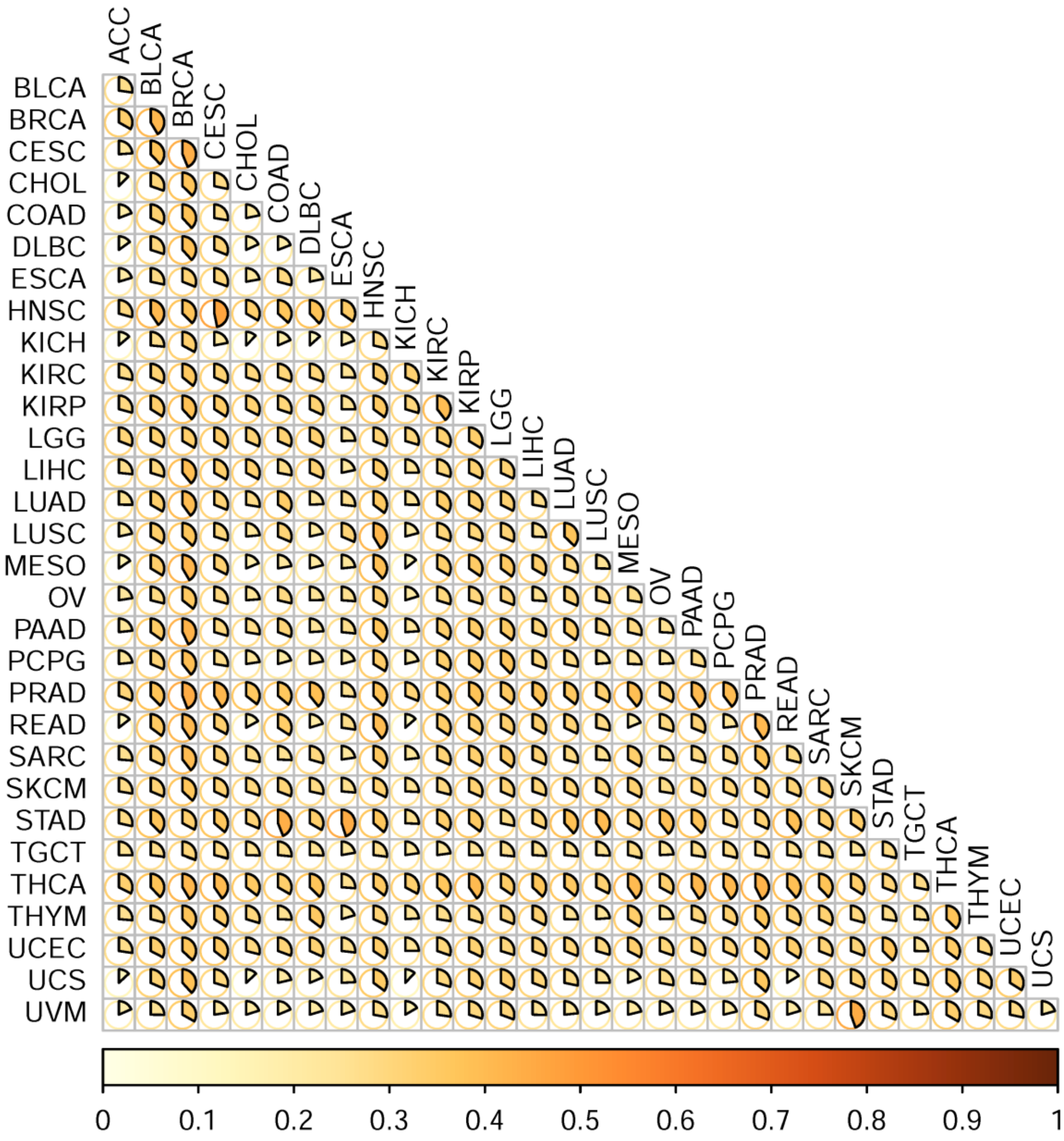


**Fig S2. Similarity matrix of ceRNA networks.** The similarity matrix shows the similarity between each pair of ceRNA networks across the 31 malignant tumors.

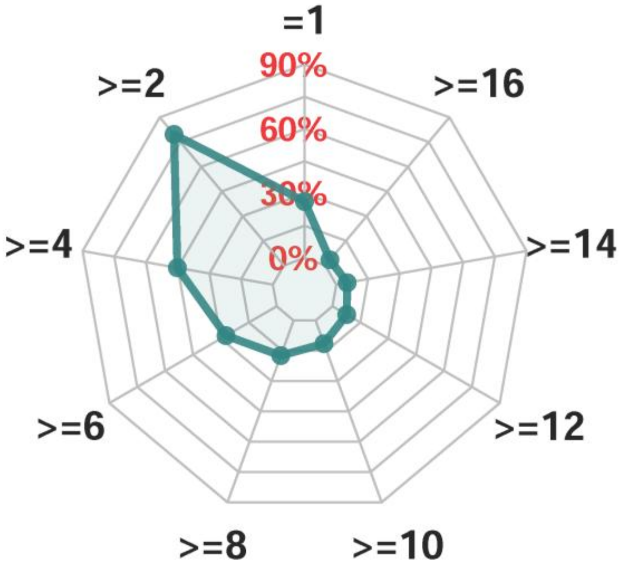


**Fig S3. The radar chart of conserved ceRNA regulation**. The radar chart shows the percentage of ceRNA interactions predicted in different number of malignant tumors.

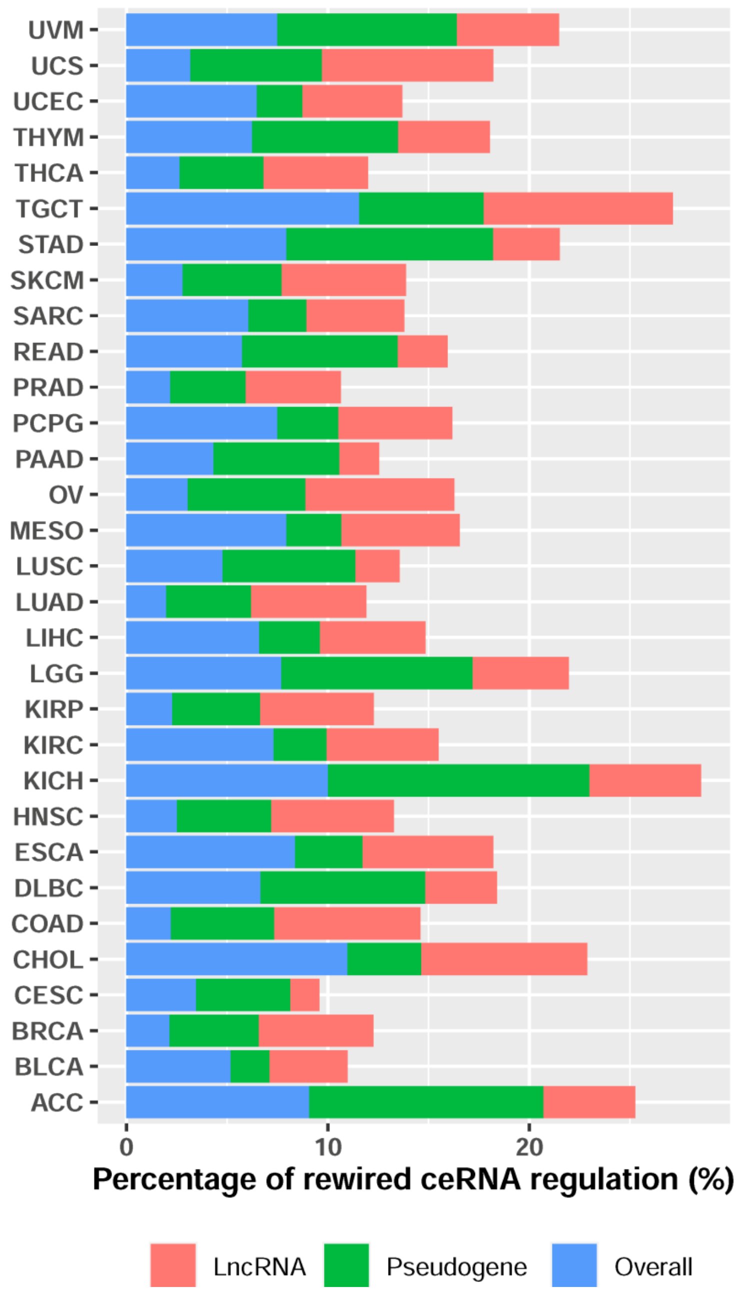


**Fig S4. Percentage of rewired ceRNA regulation.** The bar chart shows the percentage of ceRNA interactions predicted in one malignant tumor.

**a** Classification analysis of conserved ceRNA network

| Metrics | Conserved | Baseline |
|---|---|---|
| accuracy | 0.65 | 0.09 |
| average-precision | 0.77 | 0.17 |
| clp | 0.00 | 0.03 |
| coverage | 1.74 | 14.42 |
| F1 | 0.66 | 0.09 |
| hamming-loss | 0.02 | 0.06 |
| macro-AUC | 0.97 | 0.50 |
| macro-F1 | 0.60 | 0.01 |
| macro-precision | 0.65 | 0.00 |
| macro-recall | 0.59 | 0.03 |
| margin-loss | 1.74 | 14.42 |
| micro-AUC | 0.96 | 0.53 |
| micro-F1 | 0.65 | 0.09 |
| micro-precision | 0.62 | 0.09 |
| micro-recall | 0.69 | 0.09 |
| mlp | 0.06 | 0.97 |
| one-error | 0.33 | 0.91 |
| precision | 0.65 | 0.09 |
| ranking-loss | 0.06 | 0.48 |
| recall | 0.69 | 0.09 |
| subset-accuracy | 0.61 | 0.09 |
| wlp | 0.10 | 0.97 |

**b** Survival analysis of conserved ceRNA network

| Tumors | p-value | HR | HRlow95 | HRup95 |
|---|---|---|---|---|
| ACC | 5.33E-15 | 9.79 | 3.33 | 28.81 |
| BLCA | <2.22E-16 | 5.60 | 3.80 | 8.25 |
| BRCA | <2.22E-16 | 12.58 | 7.90 | 20.06 |
| CESC | <2.22E-16 | 11.78 | 6.09 | 22.78 |
| CHOL | 8.68E-11 | 11.18 | 2.23 | 55.95 |
| COAD | <2.22E-16 | 17.01 | 7.94 | 36.46 |
| DLBC | 7.13E-09 | 21.25 | 3.26 | 138.57 |
| ESCA | <2.22E-16 | 7.25 | 3.11 | 16.94 |
| HNSC | <2.22E-16 | 5.04 | 3.53 | 7.18 |
| KICH | 7.27E-06 | / | / | / |
| KIRC | <2.22E-16 | 13.69 | 6.66 | 28.16 |
| KIRP | <2.22E-16 | 17.51 | 7.28 | 42.11 |
| LGG | <2.22E-16 | 10.64 | 6.83 | 16.56 |
| LIHC | <2.22E-16 | 8.01 | 4.99 | 12.87 |
| LUAD | <2.22E-16 | 6.62 | 4.38 | 10.00 |
| LUSC | <2.22E-16 | 7.57 | 4.64 | 12.35 |
| MESO | <2.22E-16 | 7.31 | 2.99 | 17.90 |
| OV | <2.22E-16 | 5.36 | 3.51 | 8.20 |
| PAAD | <2.22E-16 | 9.48 | 4.91 | 18.31 |
| PCPG | 1.48E-04 | / | / | / |
| PRAD | 8.82E-08 | 13.02 | 3.05 | 55.55 |
| READ | 1.61E-09 | 9.29 | 2.29 | 37.63 |
| SARC | <2.22E-16 | 10.50 | 5.85 | 18.85 |
| SKCM | <2.22E-16 | 5.50 | 3.75 | 8.08 |
| STAD | <2.22E-16 | 10.01 | 6.17 | 16.24 |
| TGCT | 2.66E-03 | / | / | / |
| THCA | 8.24E-14 | / | / | / |
| THYM | 5.91E-06 | 10.52 | 1.77 | 62.59 |
| UCEC | <2.22E-16 | 20.40 | 11.37 | 36.58 |
| UCS | 1.11E-16 | 9.22 | 3.04 | 27.95 |
| UVM | <2.22E-16 | 12.86 | 3.12 | 52.96 |

**Fig S5. Analysis of conserved ceRNA regulation.** (a) Multi-class classification analysis of conserved ceRNA network in 31 malignant tumors. (b) Survival analysis of the conserved ceRNA network in 31 malignant tumors. The symbol "/" stands for an infinite value.

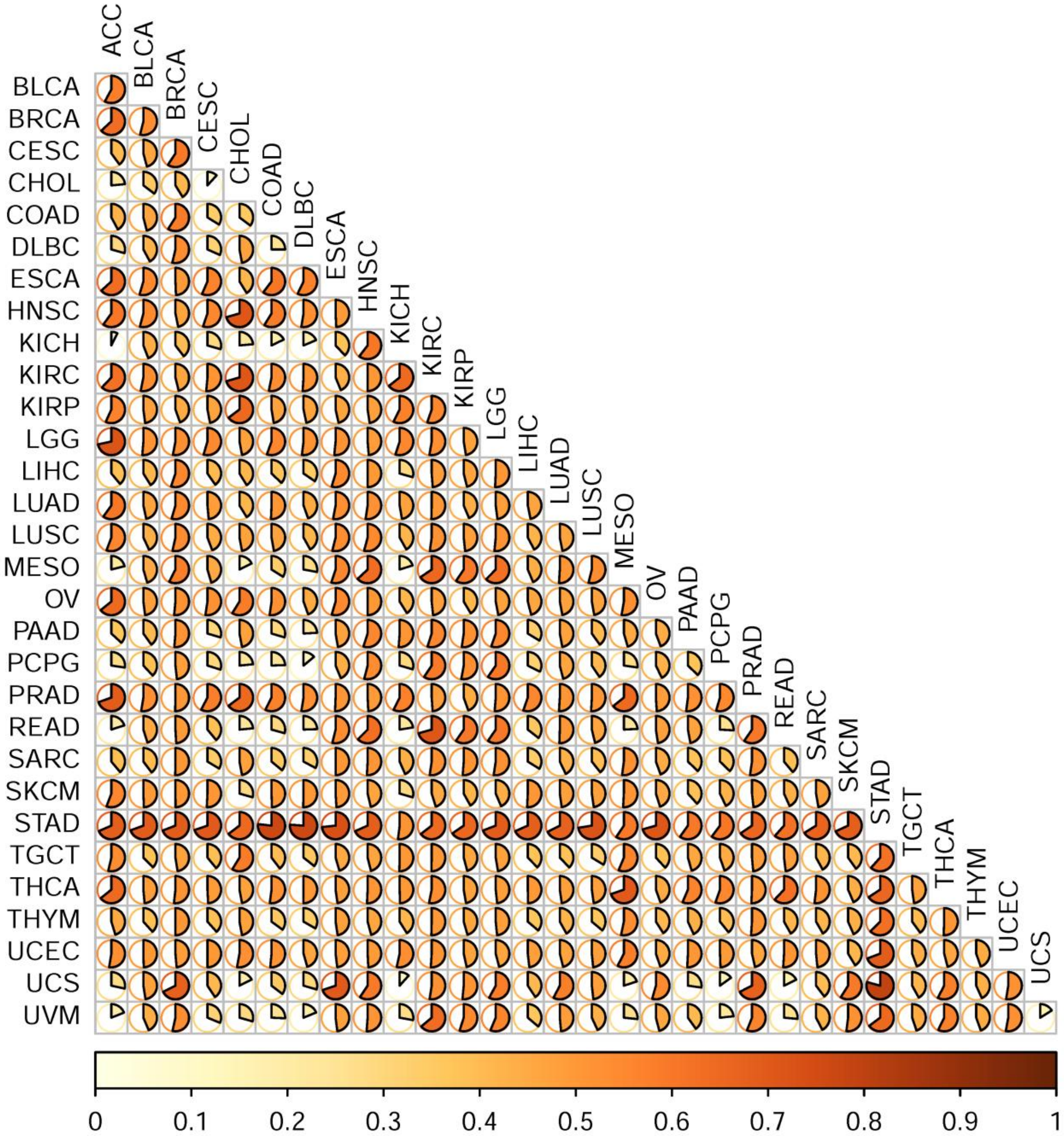


**Fig S6. Similarity matrix of hub ncRNAs.** The similarity matrix shows the similarity between each pair of hub ncRNAs across the 31 malignant tumors.

**a** Survival analysis of hub ncRNAs

| Tumors | p-value | HR | HRlow95 | HRup95 |
|---|---|---|---|---|
| ACC | 5.33E-15 | 9.79 | 3.33 | 28.81 |
| BLCA | <2.22E-16 | 12.05 | 7.43 | 19.54 |
| BRCA | <2.22E-16 | 10.72 | 5.33 | 21.55 |
| CESC | <2.22E-16 | 15.19 | 7.36 | 31.35 |
| CHOL | 1.27E-04 | 5.03 | 1.35 | 18.76 |
| COAD | <2.22E-16 | 8.23 | 3.60 | 18.82 |
| DLBC | 5.49E-08 | 14.85 | 1.97 | 111.88 |
| ESCA | <2.22E-16 | 8.82 | 3.71 | 20.97 |
| HNSC | <2.22E-16 | 14.35 | 9.20 | 22.39 |
| KICH | 6.19E-07 | / | / | / |
| KIRC | <2.22E-16 | 11.26 | 4.84 | 26.21 |
| KIRP | <2.22E-16 | 21.21 | 8.29 | 54.29 |
| LGG | <2.22E-16 | 8.16 | 4.24 | 15.69 |
| LIHC | <2.22E-16 | 13.07 | 7.60 | 22.48 |
| LUAD | <2.22E-16 | 9.84 | 5.79 | 16.72 |
| LUSC | <2.22E-16 | 12.73 | 7.24 | 22.37 |
| MESO | <2.22E-16 | 7.31 | 2.99 | 17.90 |
| OV | <2.22E-16 | 8.32 | 4.48 | 15.46 |
| PAAD | <2.22E-16 | 7.68 | 3.78 | 15.59 |
| PCPG | 5.45E-07 | / | / | / |
| PRAD | 8.68E-09 | 19.63 | 4.81 | 80.11 |
| READ | 3.53E-10 | 9.92 | 2.37 | 41.60 |
| SARC | <2.22E-16 | 8.57 | 4.45 | 16.52 |
| SKCM | <2.22E-16 | 9.45 | 5.69 | 15.70 |
| STAD | <2.22E-16 | 9.52 | 5.36 | 16.91 |
| TGCT | 2.13E-03 | / | / | / |
| THCA | 1.02E-11 | 25.53 | 7.25 | 89.86 |
| THYM | 1.08E-05 | 10.10 | 1.74 | 58.63 |
| UCEC | <2.22E-16 | 11.68 | 5.62 | 24.31 |
| UCS | 1.11E-16 | 9.17 | 2.86 | 29.37 |
| UVM | 1.11E-13 | 10.23 | 2.56 | 40.86 |

**b** Enrichment analysis of hub ncRNAs

| Tumors | #Diseases | #Cell markers | #Cancer hallmarks |
|---|---|---|---|
| ACC | 0 | 0 | 1 |
| BLCA | 68 | 7 | 7 |
| BRCA | 119 | 7 | 7 |
| CESC | 34 | 2 | 3 |
| CHOL | 0 | 1 | 0 |
| COAD | 25 | 4 | 8 |
| DLBC | 1 | 5 | 0 |
| ESCA | 35 | 6 | 6 |
| HNSC | 78 | 12 | 6 |
| KICH | 83 | 11 | 6 |
| KIRC | 142 | 17 | 7 |
| KIRP | 162 | 13 | 7 |
| LGG | 103 | 7 | 7 |
| LIHC | 154 | 4 | 7 |
| LUAD | 77 | 8 | 6 |
| LUSC | 106 | 9 | 7 |
| MESO | 52 | 7 | 7 |
| OV | 76 | 4 | 6 |
| PAAD | 130 | 14 | 7 |
| PCPG | 59 | 7 | 4 |
| PRAD | 100 | 10 | 6 |
| READ | 117 | 9 | 7 |
| SARC | 137 | 6 | 7 |
| SKCM | 56 | 6 | 6 |
| STAD | 50 | 3 | 6 |
| TGCT | 169 | 13 | 7 |
| THCA | 134 | 13 | 7 |
| THYM | 106 | 7 | 7 |
| UCEC | 152 | 9 | 7 |
| UCS | 0 | 0 | 0 |
| UVM | 167 | 9 | 7 |

**c** Classification analysis of rewired hub ncRNAs

| Metrics | Rewired | Baseline |
|---|---|---|
| accuracy | 0.93 | 0.09 |
| average-precision | 0.96 | 0.17 |
| clp | 0.00 | 0.03 |
| coverage | 0.17 | 14.42 |
| F1 | 0.93 | 0.09 |
| hamming-loss | 0.00 | 0.06 |
| macro-AUC | 0.99 | 0.50 |
| macro-F1 | 0.90 | 0.01 |
| macro-precision | 0.89 | 0.00 |
| macro-recall | 0.91 | 0.03 |
| margin-loss | 0.17 | 14.42 |
| micro-AUC | 1.00 | 0.53 |
| micro-F1 | 0.93 | 0.09 |
| micro-precision | 0.92 | 0.09 |
| micro-recall | 0.94 | 0.09 |
| mlp | 0.00 | 0.97 |
| one-error | 0.07 | 0.91 |
| precision | 0.93 | 0.09 |
| ranking-loss | 0.01 | 0.48 |
| recall | 0.94 | 0.09 |
| subset-accuracy | 0.91 | 0.09 |
| wlp | 0.00 | 0.97 |

**d** Survival analysis of rewired hub ncRNAs

| Tumors | p-value | HR | HRlow95 | HRup95 |
|---|---|---|---|---|
| ACC | 5.33E-15 | 9.79 | 3.33 | 28.81 |
| BLCA | <2.22E-16 | 13.39 | 8.28 | 21.64 |
| BRCA | <2.22E-16 | 19.60 | 10.73 | 35.78 |
| CESC | <2.22E-16 | 12.00 | 5.69 | 25.28 |
| CHOL | 8.68E-11 | 11.18 | 2.23 | 55.95 |
| COAD | <2.22E-16 | 9.29 | 3.91 | 22.09 |
| DLBC | 4.95E-09 | 21.75 | 3.28 | 144.03 |
| ESCA | <2.22E-16 | 9.28 | 3.74 | 23.05 |
| HNSC | <2.22E-16 | 10.32 | 6.41 | 16.63 |
| KICH | 7.27E-06 | / | / | / |
| KIRC | <2.22E-16 | 11.99 | 4.92 | 29.21 |
| KIRP | <2.22E-16 | 8.60 | 3.04 | 24.37 |
| LGG | <2.22E-16 | 8.52 | 4.54 | 16.02 |
| LIHC | <2.22E-16 | 14.54 | 8.59 | 24.59 |
| LUAD | <2.22E-16 | 13.16 | 7.94 | 21.81 |
| LUSC | <2.22E-16 | 9.80 | 5.31 | 18.08 |
| MESO | <2.22E-16 | 7.31 | 2.99 | 17.90 |
| OV | <2.22E-16 | 8.33 | 4.54 | 15.30 |
| PAAD | <2.22E-16 | 7.64 | 3.77 | 15.48 |
| PCPG | 3.94E-05 | 20.60 | 2.74 | 154.98 |
| PRAD | 2.60E-07 | 12.22 | 2.95 | 50.68 |
| READ | 5.28E-11 | 10.72 | 2.45 | 46.92 |
| SARC | <2.22E-16 | 13.54 | 7.42 | 24.70 |
| SKCM | <2.22E-16 | 9.47 | 5.67 | 15.80 |
| STAD | <2.22E-16 | 11.94 | 6.68 | 21.33 |
| TGCT | 3.86E-03 | / | / | / |
| THCA | 1.97E-11 | 18.58 | 4.94 | 69.94 |
| THYM | 1.02E-05 | 10.11 | 1.74 | 58.76 |
| UCEC | <2.22E-16 | 13.17 | 6.45 | 26.91 |
| UCS | 1.11E-16 | 9.17 | 2.86 | 29.37 |
| UVM | 1.89E-14 | 11.10 | 3.20 | 38.56 |

**Fig S7. Hub analysis of ncRNA synergistic competition across the 31 malignant tumors.** (a) Survival analysis of hub ncRNAs. (b) Enrichment analysis of hub ncRNAs. (c) Multi-class classification analysis of rewired hub ncRNAs. (d) Survival analysis of rewired hub ncRNAs. The symbol "/" stands for an infinite value.

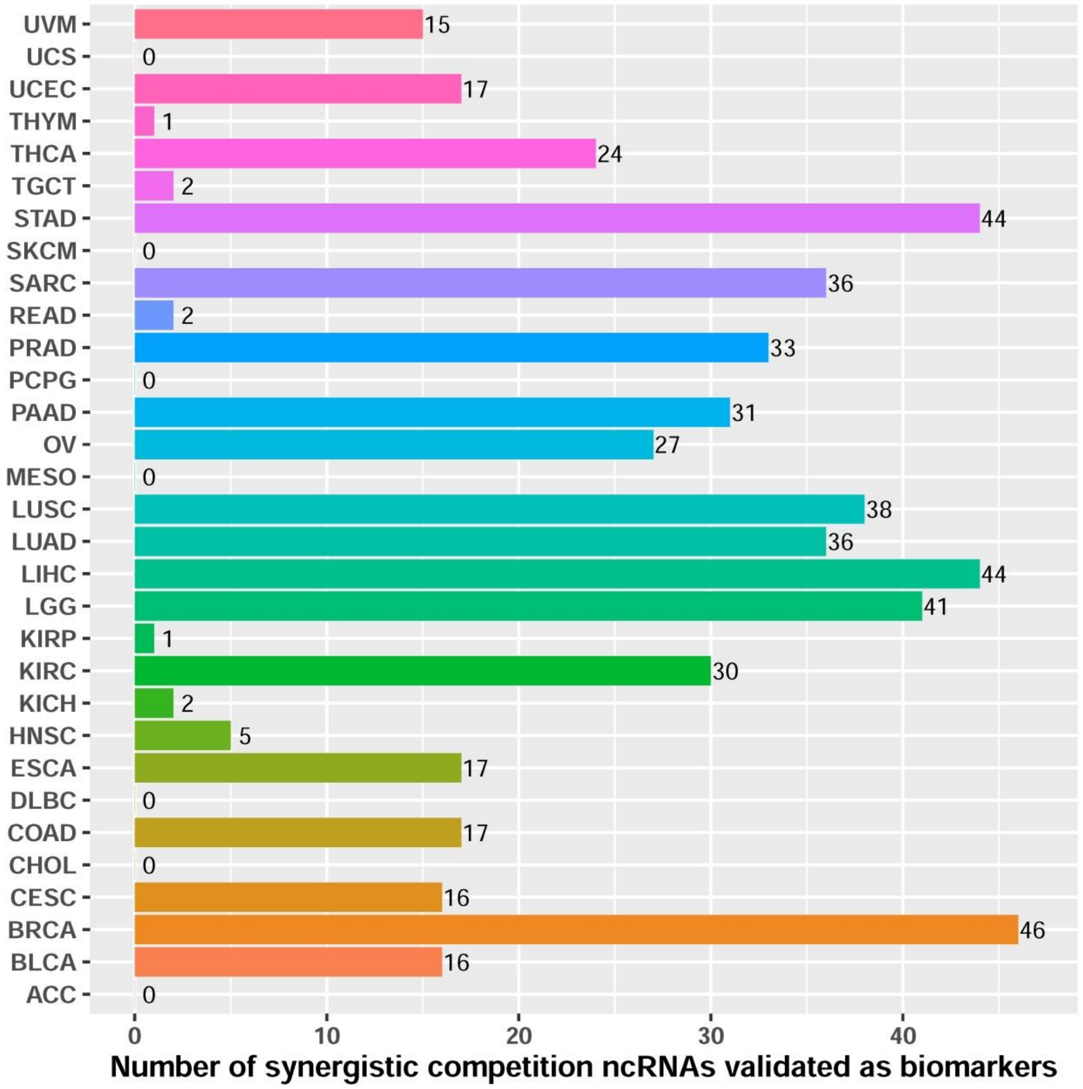


**Fig S8. Number of experimentally validated synergistic competition ncRNAs as biomarkers.** The bar chart shows the number of synergistic competition ncRNAs validated as biomarkers in the literature.
